# Analysis of justification for and gender bias in author order among those contributing equally

**DOI:** 10.1101/2024.03.01.582955

**Authors:** Ellie Rose Mattoon, Maisha Miles, Nichole A Broderick, Arturo Casadevall

## Abstract

The practice of designating two or more authors as equal contributors (EC) on a scientific publication is increasingly common as a form of sharing credit. However, EC authors are often unclearly attributed on CVs or citation engines, and it is unclear how research teams determine author order within an EC listing. In response to studies showing that male authors were more likely to be placed first in an EC listing, the American Society of Microbiology (ASM) required that authors explain the reasons for author order beginning in 2020. In this study we analyze data from over 2500 ASM publications to see how this policy affected gender bias and how research teams are making decisions on author order. Data on publications from 2018-2021 show that gender bias was largely nonsignificant both before and after authors were asked by ASM to provide an EC statement. The most likely reasons for EC order included alphabetical order, seniority, and chance, although there were differences for publications from different geographic regions. However, many research teams used unique methods in order selection, highlighting the importance of EC statements to provide clarity for readers, funding agencies, and tenure committees.

## Introduction

The number of scientific journal articles listing two or more authors as equal contributors (ECs) has steadily increased since the 1990s (1). This trend occurs concurrently with increases in the total number of authors on a publication (2) and increases in international collaboration (3). Because funding, tenure, and hiring committees often use an author’s byline position as a metric of relative contribution, EC presents a potential solution to ensure that collaborators receive the appropriate credit for their contributions (4).

Despite these efforts, EC authors still receive less credit if their name is listed second on the byline. Readers instinctively associate the first position with greater influence, prestige and importance based on the long tradition of recognizing the first author as the major contributor to published works in the biomedical sciences. This problem of misperception is accentuated by the fact that EC is often not acknowledged on reference lists or automated research profiles (4), and it is currently unclear how EC is viewed in hiring and promotions decisions (1). A 2017 survey of 6000 corresponding authors found that contribution statements were more likely to be used to assess a type of contribution rather than effort or credit (5). In addition, given the diversity of social dynamics on research teams and laboratories, the methods used to determine author order even in cases of EC vary widely from paper to paper (5).

In 2019, an analysis of 2898 scientific papers with listed EC authors found that a male author was more likely to be listed first than a female counterpart on an EC publication (6), although the phenomena was declining over time. A similar analysis of 10 journals with high-impact factors found that although there was an insignificant gender bias in basic science journals, male authors were more likely to be listed first in clinical journals (7). Women are already less likely to be credited with authorship on a project (8), and although women receive 41% of STEM doctorate degrees, they hold only 26% of STEM-related leadership roles (9). This imbalance in first author contributions parallels imbalances in citations, funding, and other forms for recognition for women scientists. Our prior findings about EC authorship led to wider conversations on how EC practices might reflect or perpetuate gender inequalities in science (10).

In response to these studies, the *Journal of Clinical Investigation* (JCI) (10) and journals of the American Society for Microbiology (ASM) (11) announced that they would be requiring corresponding authors to state the method used in assigning first-author position. In a statement, *JCI* leadership announced that they aimed to promote “discussions between authors and their supervisors that could lead to fairer choices” (10). There was also hope that by requiring explanations for the author’s order that information could help those listed second to explain the decision and obtain their fair share of credit (11). In this study we analyze the outcome of the policy requiring author order explanations among those contributing equally. We found a wide range of methods used to order EC authors, showcasing the importance of reviewing EC publications and author contributions on a case-by-case basis.

## Materials and Methods

### Data Sources

All analyzed data was taken from the submission information from 3177 publications from American Society for Microbiology journals published between 2018-2021, all of which specified two equal contributors in either the first, middle, or final position on their submission information. Starting in 2020, submissions also were asked to include an explanation for how author order was determined among equal contributors.

### Data Analysis

Methods are replicated from those used in (6). One of the coauthors manually examined each article to determine the gender of each provided author. Determination of an author’s presenting gender was primarily completed by searching an individual’s full name and affiliated institution on the Google search engine. Oftentimes, the image and pronouns on their faculty or student profile were sufficient to determine the individual’s gender presentation. For individuals who did not have an identifying photo or who had a name commonly perceived as gender-neutral (such as “Avery” or “Leslie”), that paper’s corresponding author was emailed and asked if they could provide their colleague’s gender identity. A portion of corresponding authors either did not respond after three contact attempts or asked to opt out of the study. This study was unable to account for nonbinary individuals in its analysis.

For each entry, we also recorded the byline position of the ECs (first vs. last), the country of the corresponding author, and the method of author order by the following categories: Alphabetical, Random, Other. After completing the initial analysis, an additional category, “Seniority” was added to account for the large number of entries claiming to use this method.

### Statistics

Gender bias in authorship was defined in two ways: the proportion of EC papers with two authors of different genders that presented author bylines as Male-Female, and the proportion of all EC papers that had a male in the first author spot (this includes Male-Female (M:F) entries, Male-Male (M:M) entries, and entries with three or more ECs in which a Male is listed first (M+)). For both pairings, a proportion closer to 50% was assumed to have less gender bias.

For the regional analysis of provided entries, each entry was categorized into the following world regions: North America, Europe, Asia, South America, Middle East, Africa, and Australia. Due to low sample size, South America, the Middle East, Africa, and Australia entries were either excluded from some analyses or combined into an “Other” category. Entries with corresponding authors from multiple continents were excluded from the analysis of regional characteristics of author order methods.

All data analysis was carried out using Excel.

## Results

### Summary Data Among ASM Journals

From 3172 scientific publications in 13 ASM journals published between 2019-2021, 2665 (84%) usable entries (entries for which we could obtain gender information) were identified (**Table 1**). Of the provided entries, 16% were excluded after the research team was unable to identify the ECs genders, and the corresponding author either failed to respond to email inquiries or asked to opt out of the analysis. Of the 3172 entries, 65% were successfully identified using Internet resources alone, and the success rate of email inquiries was 54% (**Table 2**).

**Table 1:**
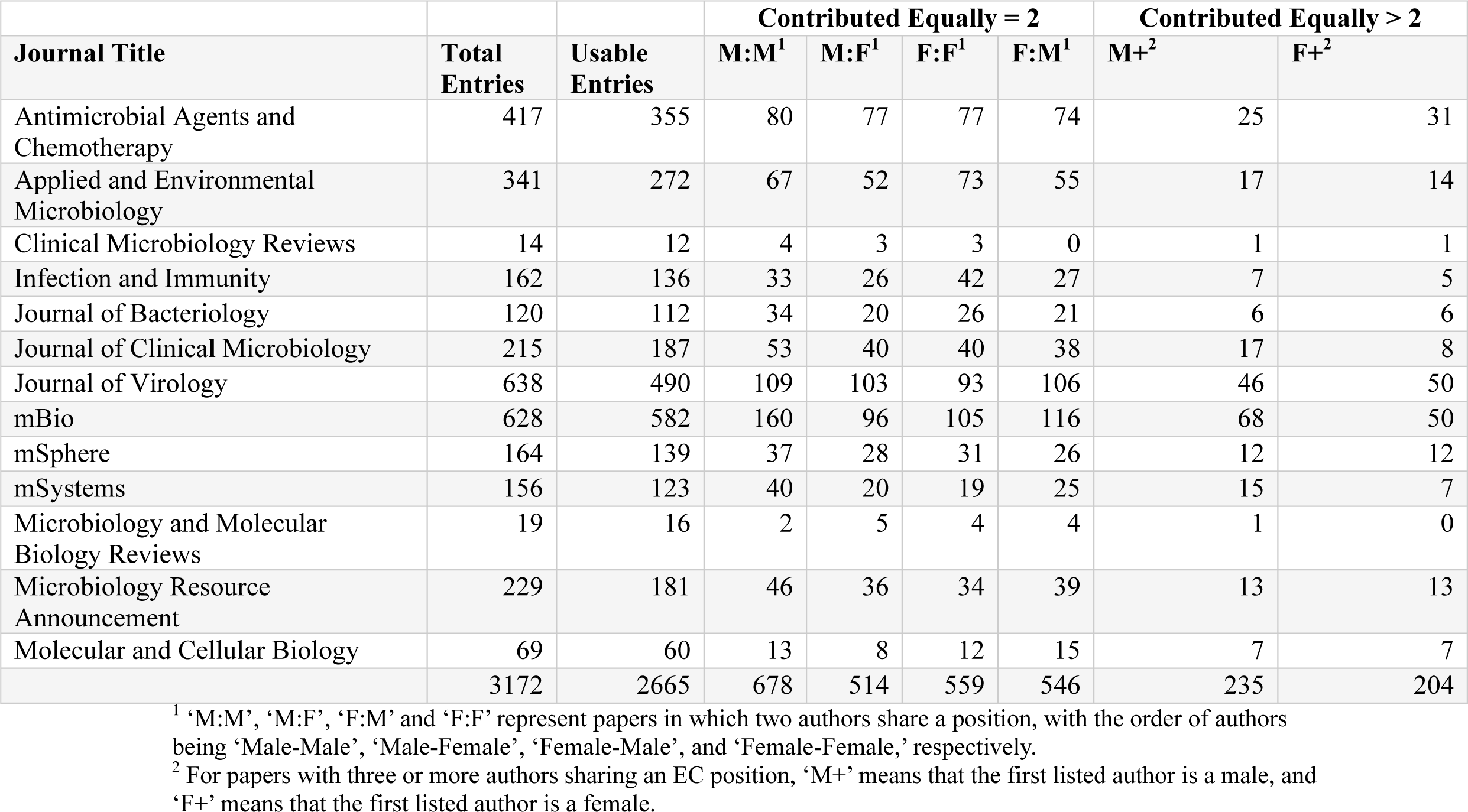
Summary data of ASM authors listed as equal contributors, organized by journal and author gender.

**Table 2:** Summary data of methods used to identify author gender (Internet vs. Email) and email inquiry success rate, organized by year of journal publication.

| **Year** | **Internet** | **Emails**<br>**(Total)** | **Successful**<br>**Emails** | **Excluded** |
| --- | --- | --- | --- | --- |
| 2019 | 684 | 385 | 197 | 188 |
| 2020 | 753 | 439 | 233 | 206 |
| 2021 | 627 | 284 | 170 | 114 |
| Total | 2064 | 1108 | 600 | 508 |
| Proportion | 0.65 | 0.35 | 0.19 | 0.16 |

### The Impact of EC justification Statement on Gender Bias

ASM began requiring EC justification statements for journals published in 2020. One entry from 2019 provided that the authors were ordered alphabetically without being asked to (**Table 3**).

**Table 3:** EC author gender, byline order, and method of determining first author organized by year. Submissions in 2019 were not asked to provide reasons for why a given author was included first on the byline.

| Year | M:M<sup>1</sup> | M:F<sup>1</sup> | F:M<sup>1</sup> | F:F<sup>1</sup> | M+<sup>2</sup> | F+<sup>2</sup> | First | Last | Alphabetical | Other | Random | Seniority |
| --- | --- | --- | --- | --- | --- | --- | --- | --- | --- | --- | --- | --- |
| 2019 | 229 | 159 | 192 | 184 | 69 | 72 | 745 | 124 | 1 | 0 | 0 | 0 |
| 2020 | 251 | 193 | 203 | 209 | 83 | 76 | 836 | 150 | 253 | 338 | 62 | 228 |
| 2021 | 198 | 162 | 151 | 166 | 83 | 56 | 656 | 124 | 185 | 276 | 68 | 160 |
| Sum | 678 | 514 | 546 | 559 | 235 | 204 | 2237 | 398 | 439 | 614 | 130 | 391 |

<sup>1</sup> ‘M:M’, ‘M:F’, ‘F:M’ and ‘F:F’ represent papers in which two authors share a position, with the order of authors being ‘Male-Male’, ‘Male-Female’, ‘Female-Male’, and ‘Female-Female,’ respectively.
<sup>2</sup> For papers with three or more authors sharing an EC position, ‘M+’ means that the first listed author is a male, and ‘F+’ means that the first listed author is a female.

Approximately nine percent of entries from 2020 and 2021 explained that their EC authors contributed equally, but they failed to add any supplementary information as to why a given author was added first, despite the journal requirements, which could reflect inadequate enforcement.

Other factors are important to note when examining the below table. Several 2020 and 2021 statements gave multiple reasons for author order (for example, alphabetical order and seniority) and thus are coded into multiple categories (**Table 3**). Some entries provided multiple EC statements for both the first and last author positions and are coded into both the “First” and “Last” category, while other entries only had EC authors in a middle byline position and thus are coded into neither category (**Table 3**).

When defining gender bias as any male author placed first, there was a nonsignificant difference between entries submitting before and after ASM’s policy change (p=.23). This pattern continued when looking only at M:F papers in contrast to F:M papers (p=.16). Previous research has suggested that the phenomenon of listing male authors first had been decreasing over time (6), with submissions with a male first author approaching 50% in 2019 (**Figure 1**). However, by 2021 male-first entries approached 54%. This increase did not achieve significance at the 0.05 level (Chi square, p=.12), but the finding suggests the need for future monitoring of EC ratios.

**Figure 1:**
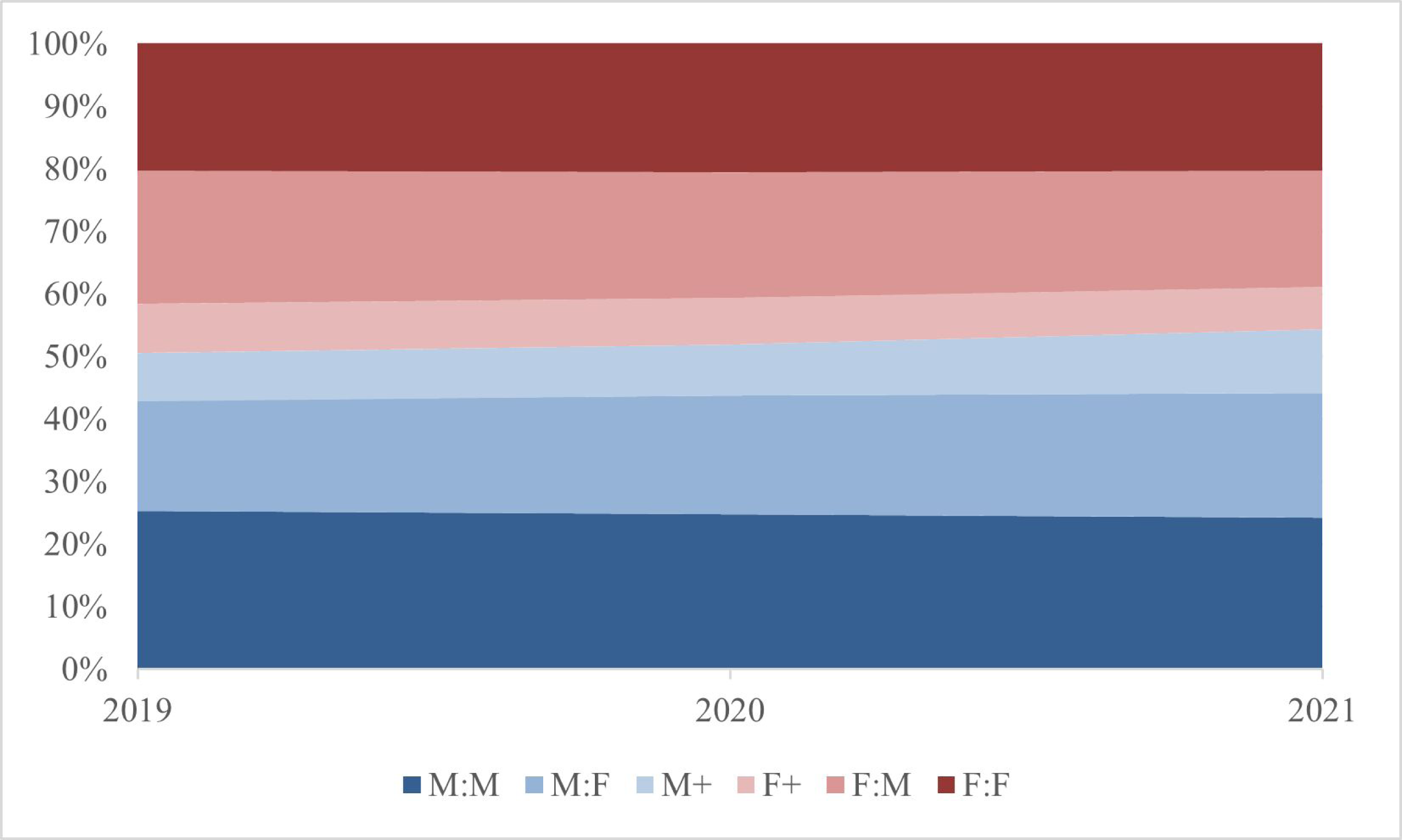
Proportion of gender combinations among EC authors in ASM journals between 2019- 2021. ‘M:M’, ‘M:F’, ‘F:M’ and ‘F:F’ represent papers in which two authors share a position, with the order of authors being ‘Male-Male’, ‘Male-Female’, ‘Female-Male’, and ‘Female- Female,’ respectively. For papers with three or more authors sharing an EC position, ‘M+’ means that the first listed author is a male, and ‘F+’ means that the first listed author is a female. The proportion of combinations in which a male author is listed first is shown in various shades of blue, while female-first combinations are shown in shades of red.

When viewing data from all three years of entries, a male author was listed first 53% of the time (**Figure 2**). While 20% of entries were Female-Male, 19% of entries were Male-Female. This data suggests that, at least in ASM journals, gender bias among EC author order was minimal between 2019-2021.

**Figure 2:**
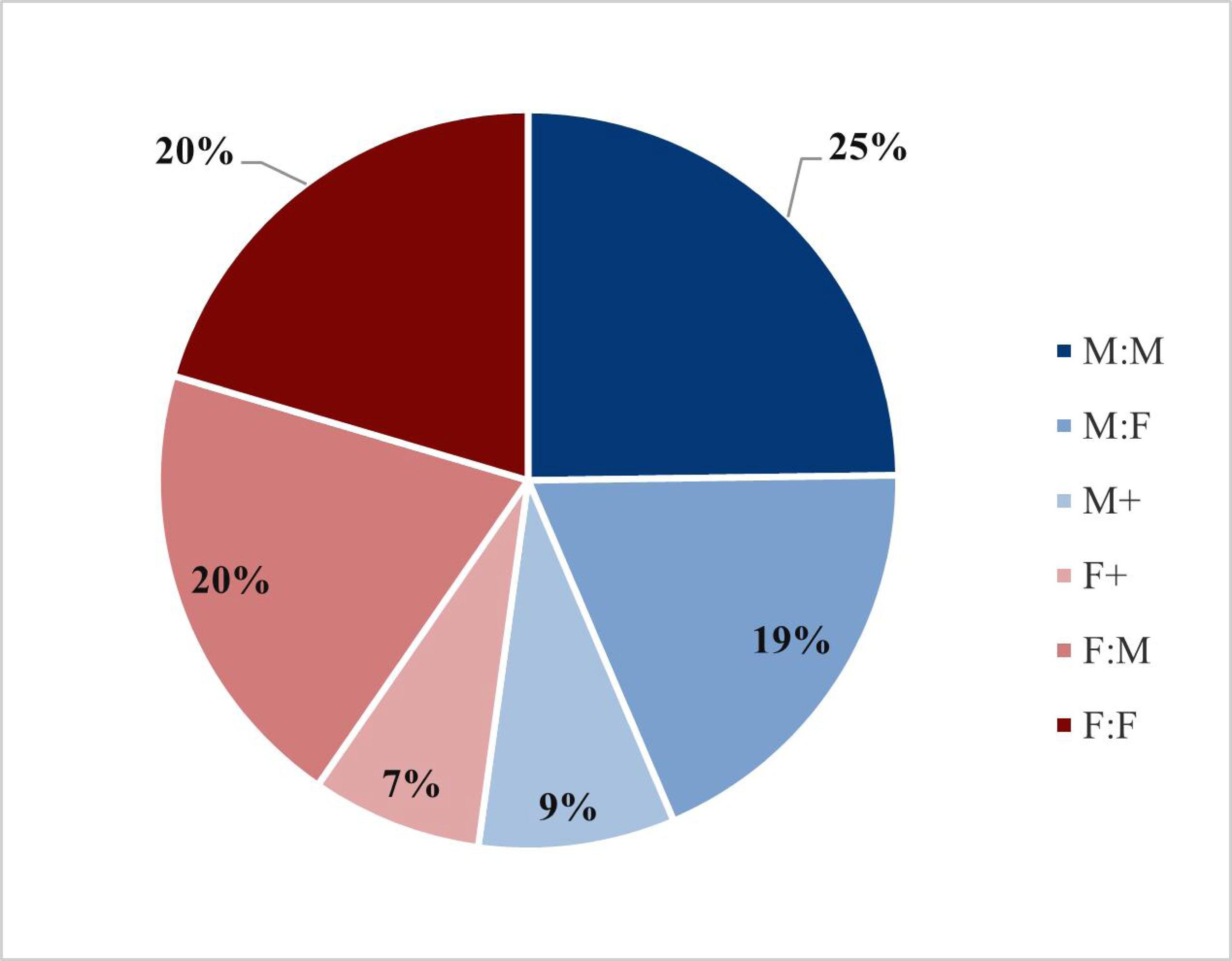
Proportion of gender combinations among EC authors in all ASM journals between 2019-2021. ‘M:M’, ‘M:F’, ‘F:M’ and ‘F:F’ represent papers in which two authors share a position, with the order of authors being ‘Male-Male’, ‘Male-Female’, ‘Female-Male’, and ‘Female-Female,’ respectively. For papers with three or more authors sharing an EC position, ‘M+’ means that the first listed author is a male, and ‘F+’ means that the first listed author is a female. The proportion of combinations in which a male author is listed first is shown in various shades of blue, while female-first combinations are shown in shades of red.

### Given Reasons for Author Order

Corresponding authors gave a variety of reasons for their EC author order, with most entries being ordered alphabetically, randomly (such as by a coin toss or drawing straws), or by seniority (**Figure 3a**). Among these three methods, alphabetical order was the most commonly used method (**Figure 3b**). Even among these methods, authors diverged in their use of forward or reverse alphabetical order and forward or reverse seniority. With regards to the methods used for ordering authors, the two-author EC entries that listed a female first were significantly less likely than male-first entries to employ chance as an ordering mechanism (p=.047).

**Figure 3:**
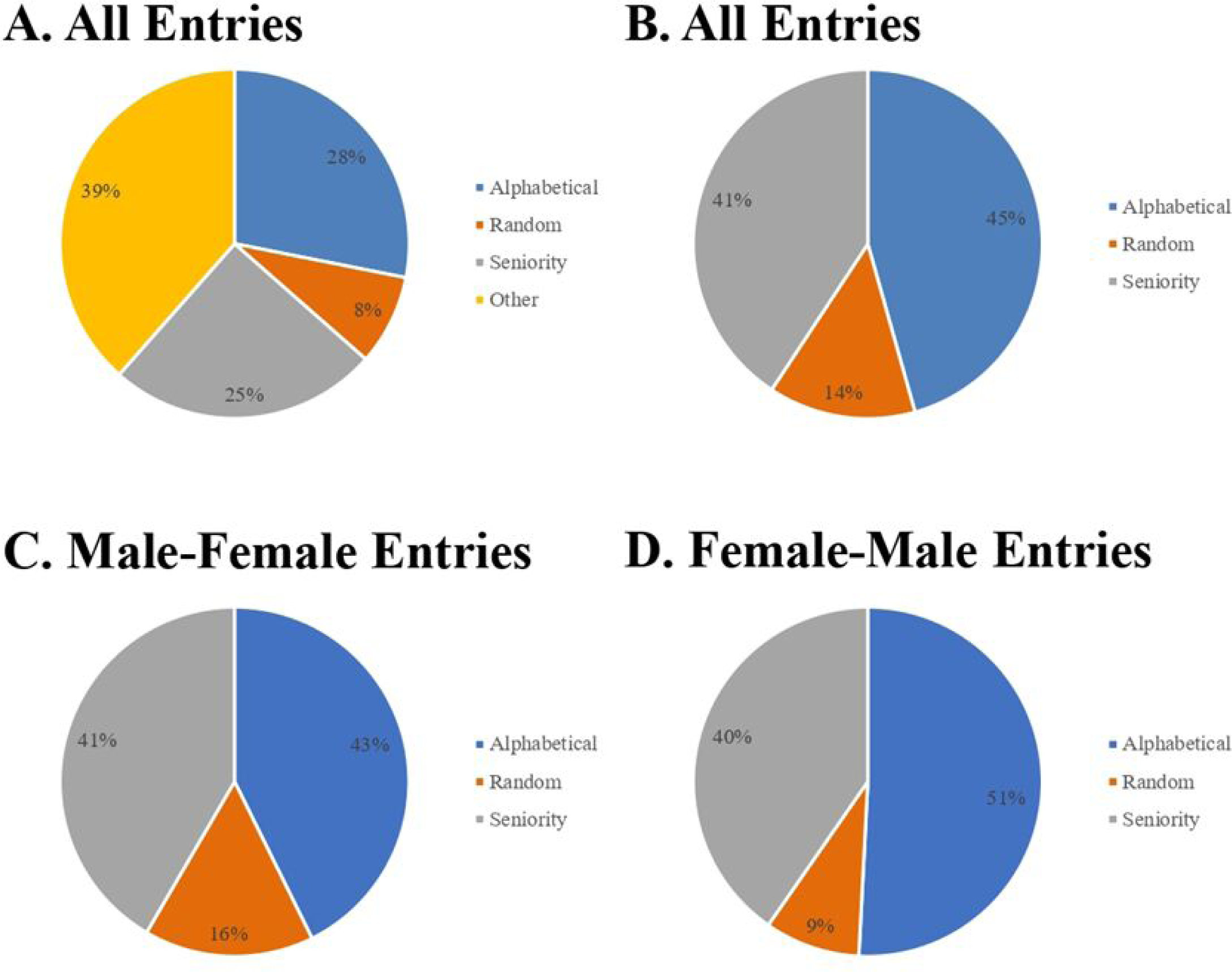
Author-provided justification for ordering EC authors among entries which provided an answer. (a) Percentage of cited methods between alphabetical, random, seniority, or other categories. Some entries listed more than one method (n=1561). (b) Percentage of cited methods between alphabetical, random, and seniority categories. Some entries listed more than one method and thus are coded into multiple categories (n=960). (c) Percentage of cited methods between alphabetical, random, and seniority categories among entries with a Male-Female EC byline (n=178). (d) Percentage of cited methods between alphabetical, random, and seniority categories among entries with a Female-Male EC byline (n=181).

A significant portion of entries used other methods such as mutual agreement, which collaborator started or finished the project, or which collaborator took on most manuscript-writing responsibilities as opposed to raw research (or vice versa). For 163 entries there was no reason for EC author order, which may reflect a lack of clarity in submission instructions or lack of journal enforcement of this requirement.

Several entries gave more unique explanations for how they determine EC author order and were also categorized as ‘other’. These explanations to include a video game (12), first name length (13), or a collaborator’s personal connection to the manuscript topic (14). One entry explicitly stated that it prioritized the female collaborator (15). Multiple entries stated that they based their order on stroke count of the author’s Chinese names (16–20), a practice used in China to organize names in a uniform order.

From these methods, the research team investigated whether certain methods were associated with a higher likelihood of gender bias. For example, since women tend to occupy fewer STEM leadership positions (9), they may be less likely to be listed first if a lab orders authors based on seniority. While all methods resulted in a male gender bias close to 50%, male authors were more likely to be listed first among all EC categories either if seniority was used to order authors or if an explanation for EC order was not provided (**Figure 4a-b**).

**Figure 4:**
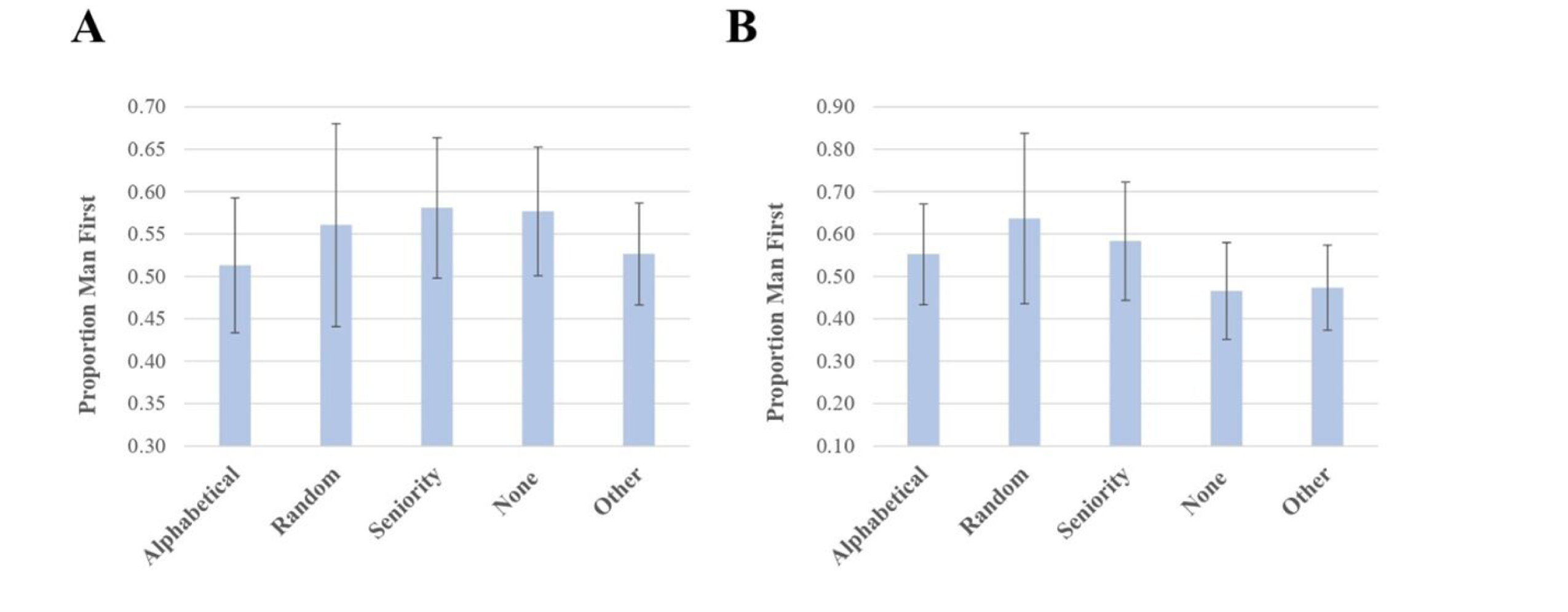
Gender bias in EC entries organized by method used to determine author order. Bars represent 95% Confidence Interval. Entries that cited multiple methods or that had multiple EC author positions were excluded. (a) Percentage of all EC entries with a male first author (n=719) (b) Percentage of two-author gender discordant EC entries that list the male collaborator first (n=305).

In past studies, researchers have suggested that using mutual agreement to determine author order disadvantages women, whom may be less likely to negotiate (21) or promote their accomplishments (22). Our data had twenty-one entries that listed the phrase “mutual agreement” in their explanation, eleven of which listed a male author first and nine of which listed a female author first. This sample is too small to support or refuse the above hypothesis.

### Regional Variations in Results

The publications our team analyzed came from research groups based in seven different world regions: North America, South America, Europe, Africa, the Middle East, Asia, and Australia. The majority of entries came from North America, Europe, and Asia. Gender bias did not differ significantly between world regions (**Figure 5a-d**).

**Figure 5:**
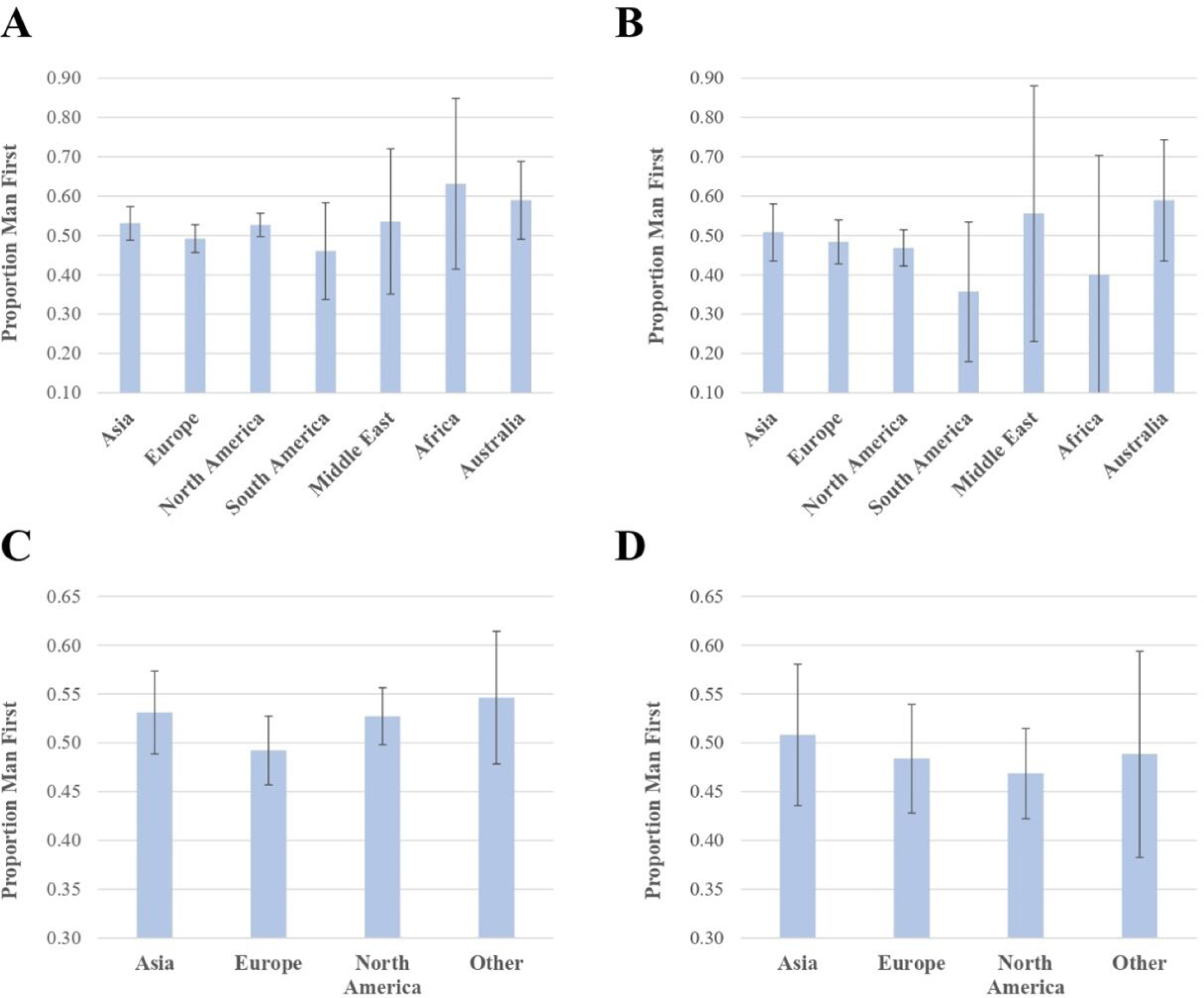
Male gender bias among EC authors organized by world region. Percentages closer to 50% are assumed to have less overall gender bias. Bars represent 95% confidence interval. Publications that were the result of inter-continental collaborations were excluded, and one publication can have multiple EC positions. (a) Percentage of all EC publications in which a male author is listed first by world region (n=2629). (b) Percentage of two-author gender- discordant EC publications in which a male is listed first (n=1027). (c) Percentage of all EC publications in which a male author is listed first by Europe, North America, Asia, and other regions (n=2629). (d) Percentage of two-author gender-discordant EC publications in which a male is listed first (n=1027).

Among Europe, North America, and Asia, regional patterns emerged in terms of how EC authors were ordered (**Figure 6**). For example, research teams in Asia were significantly less likely to alphabetize their authors (p< 0.00001) and more likely to use either seniority (p=.002) or chance (p=.00002) compared to European and North American research teams. When comparing methods between European and North American research teams, no statistically significant differences were determined.

**Figure 6:**
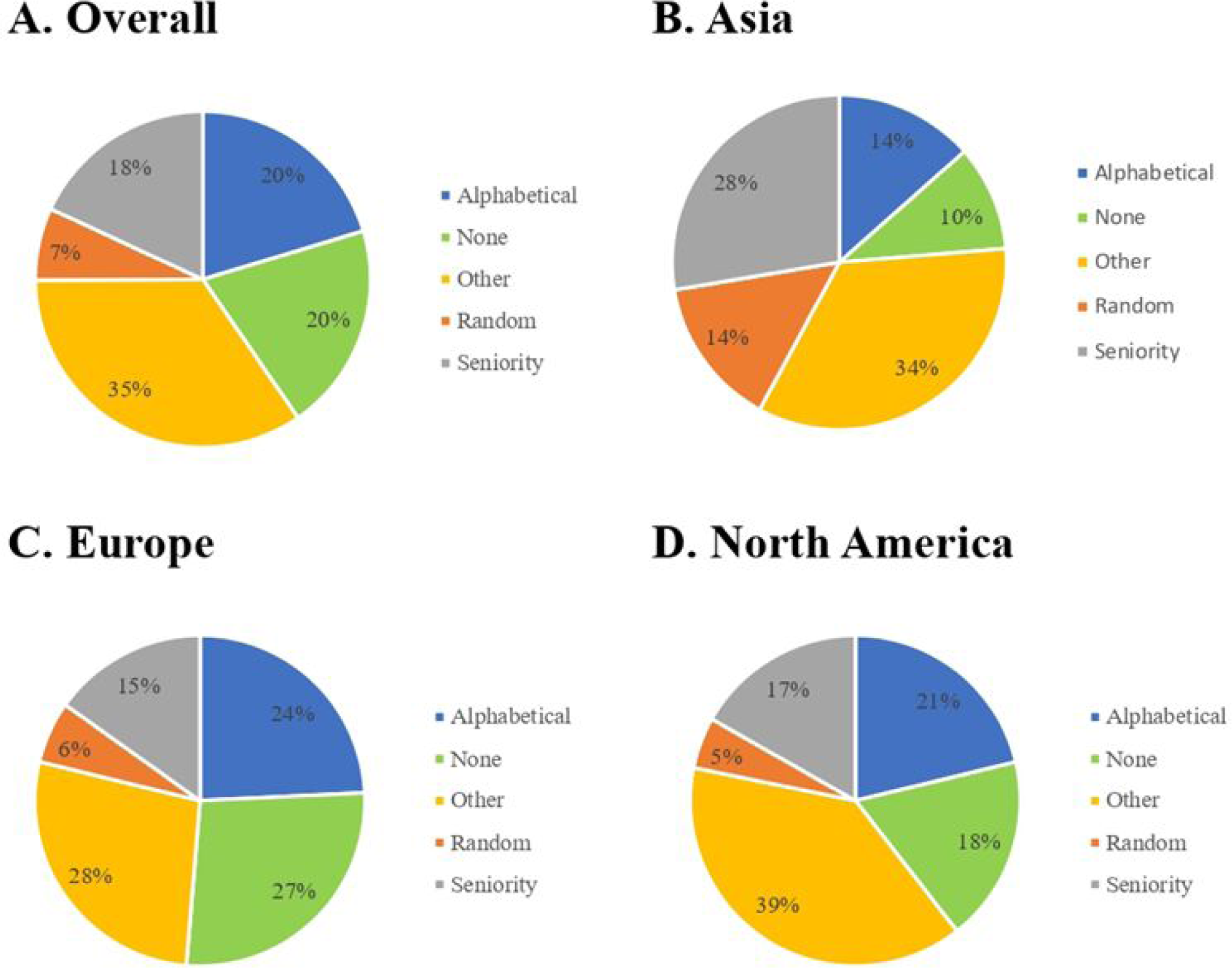
Distribution of EC ordering methods by Europe, Asia, and North America. Publications that were the result of inter-continental collaborations or that cited multiple methods of ordering authors were excluded. (a) Distribution of methods in all three regions, including the “Other or “None” category (n=1591). (b) Distribution of methods used in papers with research teams based in Asia, including the “Other” or “None” category (n=304). (c) Distribution of methods used in papers with research teams based in Europe, including the “Other” or “None” category (n=462). (d) Distribution of methods used in papers with research teams based in North America, including the “Other” or “None” category (n=697).

## Discussion

In this study, we analyzed the gender distribution and byline order of EC authors before and after ASM began requiring corresponding authors to explain how bylines were determined. Past studies had shown that male authors were in some instances more likely to be placed first in an EC byline due to the prestige associated with being a first author (6). Although this pattern of gender bias has been on the decline (6), determining more recent EC byline practices among research groups is important in assessing the best policies to ensure scientists receive fair attribution for their work. In this regard, we noted a slight but noticeable increase in publications with males in the first author position after the requirement of written explanations in 2020.

Although this difference did not reach statistical significance (Chi Square p=.012), it supports several other studies which note a general decline in female first-authorship across scientific fields in 2020 due to the COVID-19 pandemic (23,24). The pandemic disproportionately affected female scientists due to difficulties securing childcare (23), and it’s possible that our observed trend was a product of this phenomena. Nevertheless, continued monitoring is warranted as it would be unfortunate if a requirement intended to increase fairness inadvertently introduced a new bias against women. Overall, data on how research teams determine EC order provides an important glimpse into the understudied diversity of laboratory social dynamics (25).

Every EC author pair, no matter the research group a publication comes from, is presented the same way at the top of a journal article. However, every author pair is different and different laboratories appear to use different methods for determining which author is placed first in an EC pair. The results show that while most research groups use some combination of chance, alphabetization, or seniority to determine author order, even these methods are hardly a monolith. For example, some laboratories might use seniority to prioritize an author who has been in the lab for longer, while other laboratories might use reverse seniority to ensure an untenured author gets a more prestigious entry on their CV. Some laboratories base their decisions on other factors like who writes a manuscript, who finishes a project, or even who wins a game. This diversity highlights the importance of providing an EC statement so readers can have an accurate picture of how the author order was determined.

Specific methods of ordering EC authors often had a close to 50% chance of listing either a man or a woman first. However, our analysis did find that Male-Female entries were significantly more likely than Female-Male entries to employ chance. It is difficult to interpret this finding, as chance is often thought of as the method that would result in the least amount of bias. It’s possible that, as less than 10% of all the entries analyzed employed chance, this phenomenon would not appear in a larger sample size.

Overall, most world regions display a diversity of methods to determine author order, but research teams based in Asia are less likely to use alphabetical order for their EC authors. One possible explanation for this finding is that research teams in Asia are more likely to use native writing systems rather than English or a western language amenable for name ordering using the Latin alphabet. Indeed, several research groups based in China reported using stroke count of the researchers’ Chinese names to properly organize them. Given the increasing international nature of scientific publishing, journals should be conscious of this and other cultural differences between global research teams when recommending more “objective” methods of standardizing author order.

This study is not without limitations. To start, the data was analyzed using a man/woman binary and did not collect data on researchers who identify as non-binary. Requesting voluntary gender identity information upon submission or employing a larger sample size could help determine if these researchers experience gender as in EC deliberations. Additionally, an author’s gender was often identified based solely on the author’s name. While further investigation was conducted for authors with names that were determined to be gender-neutral, there is a possibility that authors with seemingly gender specific names could have been misidentified.

The data set was limited to published articles in a collection of journals focused on microbiology and allied sciences, so we do not know its applicability to other fields. Alphabetical ordering is a much more established convention in the fields of mathematics, economics, physics, and business, so results might skew towards this method if this study were replicated in these fields (26). In addition, even when some authors were asked to elaborate on how they determined byline order, nine percent of published papers did not include an explanation. This may reflect a lack of clarity in the submission instructions, which could be improved in future analyses through enforcement of journal requirements.

Even if bias for placing male EC authors first has largely declined, its legacy likely remains, particularly when it comes to the CVs of senior female scientists who began their careers when this bias was more prominent. Getting many first-author publications is seen as important for establishing young scientists’ careers, and this pattern could contribute to current inequities such as the fact that women make up 60.81% of research staff but 34.82% of faculty members (8).

More research is needed in this area to determine what the legacy of gender bias from previous decades has on women’s careers.

## Acknowledgements

There was no specific funding for this study. E.R.M. is supported in part by RO1 AI052733.

N.A.B. is supported in part by R35GM128871. K.M.S. is supported in part by NIH T32GM136577 and F30AG077736. A.C. is supported in part by R01 AI152078, R01 HL059842, and AI052733.

